# Waveform detection by deep learning reveals multi-area spindles that are selectively modulated by memory load

**DOI:** 10.1101/2021.05.14.444188

**Authors:** Maryam H. Mofrad, Greydon Gilmore, Seyed M. Mirsattari, Jorge G. Burneo, David A. Steven, Ali Khan, Ana Suller Marti, Lyle Muller

## Abstract

Sleep is generally considered to be a state of large-scale synchrony across thalamus and neocortex; however, recent work has challenged this idea by reporting isolated sleep rhythms such as slow-oscillations and spindles. What is the spatial scale of sleep rhythms? To answer this question, we adapted deep learning algorithms initially developed for detecting earthquakes and gravitational waves in high-noise settings for analysis of neural recordings in sleep. We then studied sleep spindles in non-human primate ECoG, human EEG, and clinical intracranial recordings (iEEG) in the human. We find a widespread extent of spindles, which has direct implications for the spatiotemporal dynamics we have previously studied in spindle oscillations (Muller et al., 2016) and the distribution of memory engrams in the primate.

---

Consolidation of long-term memories requires precise coordination of pre- and post-synaptic spikes across neocortex. New memories are transferred from hippocampus to neocortex for long-term storage (McClelland et al., 1995; Rasch and Born, 2007), where interconnections within a sparse, distributed neuron group are strengthened until their activity becomes hippocampus-independent (Frankland and Bontempi, 2005). Computational studies have identified neural oscillations as a potential mechanism to regulate synaptic plasticity (Masquelier et al., 2009; Song et al., 2000) and create precise spike timing (Cassenaer and Laurent, 2007; Muller et al., 2011). Further, experiments have shown that the stage 2 sleep “spindle” oscillation influences spiking activity (Contreras and Steriade, 1995; Kandel and Buzsáki, 1997; Peyrache et al., 2011) and causally contributes to sleep-dependent consolidation of long-term memory (Mednick et al., 2013). It remains unclear, however, precisely how this rhythm can coordinate activity across areas in neocortex for synaptic plasticity and long-term storage to occur.

While early recordings in anesthetized animals (Andersen et al., 1967; Contreras et al., 1996) and human EEG (Achermann and Borbély, 1998) indicated that sleep spindles generally occur across a wide area in cortex, creating a state of large-scale synchrony (Sejnowski and Destexhe, 2000; Steriade, 2003), recent work in intracranial recordings from human clinical patients has challenged this idea by reporting isolated, “local” sleep spindles (Andrillon et al., 2011; Nir et al., 2011; Piantoni et al., 2016; Sarasso et al., 2014, but see Frauscher et al., 2015). Because spindles are intrinsically related to sleep-dependent consolidation of long-term memory (Clemens et al., 2005; Gais et al., 2002; Mednick et al., 2013), this difference in reported spatial extent of the spindle raises an important question for the organization of engrams established through sleep-dependent memory consolidation. Recent evidence using *cFos* mapping in animal models suggests these engrams are distributed widely across brain areas (Kitamura et al., 2017; Roy et al., n.d.), which is consistent with previous imaging evidence in the human (Brodt and Gais, 2020; Wheeler et al., 2000). Taking these points together, we reasoned that widespread, multi-area spindles may occur more often than previously reported in primate and human cortex. If this were the case, these widespread spindles could provide the mechanism needed to link populations distributed widely across the cortex for sleep-dependent memory consolidation.

Plasticity of long-range excitatory connections linking distant neuron groups occurs through spike-time dependent plasticity (STDP) (Bi and Poo, 1998; Markram et al., 1997), for which presynaptic vesicle release and postsynaptic spiking must occur with a precision of a few milliseconds (Magee and Johnston, 1997). Long-range synaptic connections in cortex result primarily from excitatory pyramidal neurons (Schüz and Braitenberg, 2002; Sholl, 1956), and these connections could provide a link among local networks representing pieces of a memory in different brain regions. The key missing piece is to understand how spindles can guide specific long-range excitatory connections to strengthen during sleep-dependent memory consolidation. We thus hypothesized that widespread, multi-area spindles might provide this mechanism.

Reliably detecting individual spindles in noisy sleep recordings, however, is challenging. Spindle oscillation amplitudes differ across regions in cortex (Frauscher et al., 2015). Furthermore, oscillation amplitudes may differ significantly across recording sites simply due to variation in electrode properties (Kappenman and Luck, 2010; Nelson and Pouget, 2010). For these reasons, we reasoned that fixed amplitude thresholds, which are a technique common across methods for spindle detection, may only detect the largest amplitude events, potentially leading to an underestimation of spatial extent. To address this question, we adapted deep learning algorithms initially developed for detecting earthquakes (Perol et al., 2018) and gravitational waves (George and Huerta, 2018) in high-noise settings to analysis of neural recordings in sleep. We studied sleep spindles in macaque non-human primate (NHP) electrocorticogram (ECoG), human electroencephalogram (EEG), and, finally, clinical intracranial electroencephalogram (iEEG) recordings, which provide a window into the circuits of the human brain at one of the highest spatial resolutions possible (Lachaux et al., 2012; Mukamel and Fried, 2012). Our approach, which detects a range of clearly formed large- and small-amplitude spindles during sleep, reveals that the spatial extent of spindles, defined here in terms of co-occurrence across electrode sites, is widely distributed over a broad range of cortex. In particular, multi-area spindles are much more frequent than previously estimated by amplitude-thresholding approaches, which tend to select only the highest-amplitude spindles and could miss events that transiently fall below threshold. Importantly, these results were additionally verified using a signal-to-noise ratio (SNR) approach (Muller et al., 2016), which is a conservative but approximately amplitude-invariant technique closely related to the constant false alarm rate (CFAR) method used in radar (Richards, 2005). Lastly, in human sleep EEG after low- and high-load visual working memory tasks, our method detects an increase in regional and multi-area spindles uniquely following a high-load visual memory task. Taken together, these results provide substantial evidence of a specific role for spindles in linking neuron groups distributed widely across cortex during memory consolidation.

Sleep recordings from both human and NHP were obtained from electrodes ranging from the traditional scalp EEG to invasive intracranial EEG electrodes (Figure 1a). To verify the quality of spindles detected by our convolutional neural network (CNN) model (Figure 1c), we first compute average power spectral densities (PSDs) over spindle and non-spindle windows. The average PSD of detected spindle events shows an increase in the 11-15 Hz spindle frequency range (red lines, Figure 1b), while non-spindle events do not show a corresponding increase (black lines, Figure 1b). Spindles detected by the CNN are well-formed, consistent with standard morphology (Loomis et al., 1935; Newton Harvey et al., 1937; Silber et al., 2007) (Figure 1d), and in agreement with previously observed durations (0.69 ± 0.004 seconds, NHP ECoG; 0.87 ± 0.006 seconds, EEG; 0.74 ± 0.003 seconds, iEEG) (Fernandez and Lüthi, 2020; Takeuchi et al., 2016; Warby et al., 2014). To further validate spindles detected by the CNN, we designed a time-shifted averaging approach for application to recordings with only a 1 Hz highpass filter applied (thus excluding any potential effects from lowpass filtering). To do this, we collected signals from detected spindles, filtered at a 1 Hz highpass, time-aligned the events to the largest positive value within the detected window (corresponding to a positive oscillation peak), and then computed the average across aligned events. With this approach, the average over detected spindles exhibited clear 11-15 Hz oscillatory structure (black line, Supplementary Figure 1), while no oscillatory structure is observed when averaging over time-matched randomly selected non-spindle activity (dashed red line, Supplementary Figure 1). This result demonstrates that spindles detected by the CNN exhibit the correct structure even in a mostly raw, unprocessed signal with no lowpass filtering applied. Finally, we compared the amplitude distribution of spindles detected by the CNN and amplitude-thresholding (AT) approach. In the intracranial recordings (ECoG and iEEG), AT detects a subset of spindles that are significantly higher-amplitude than those detected by the CNN (p < 0.02, NHP ECoG recordings; p < 1 x 10^−14^; iEEG recordings, one-sided Wilcoxon signed-rank test; n.s. in EEG), consistent with the expectation that AT will select the largest amplitude events. The CNN, however, can find well-formed spindles that are both large and small in amplitude (Supplementary Figure 2). This improved resolution allows us to study the spatial extent of spindles in an approximately amplitude-invariant manner.

**Figure 1.**
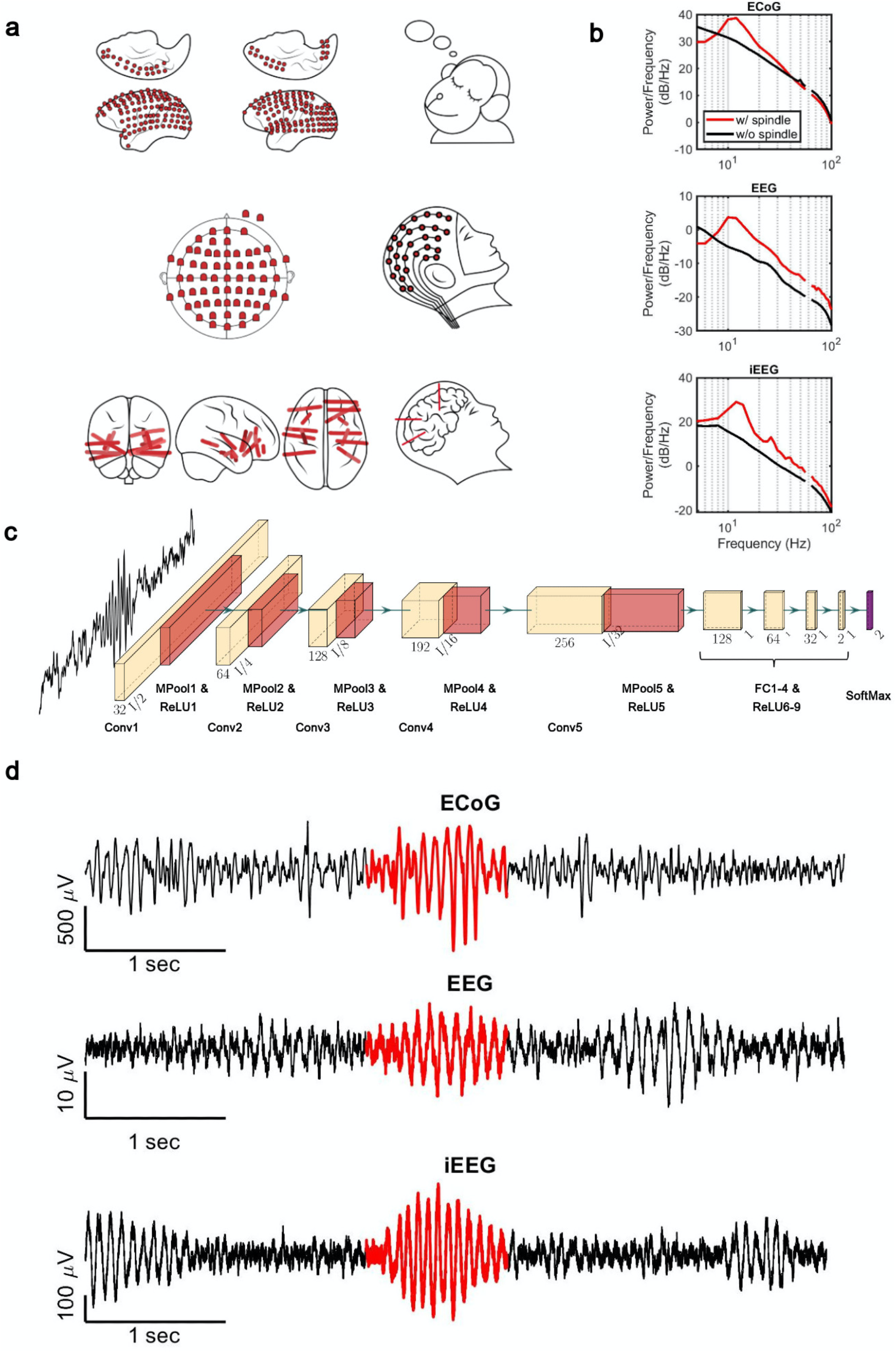
**(a)** Electrode placement of multichannel ECoG recordings of two macaques **(top)**, high-density scalp EEG used for recordings after high and low visual memory tasks **(middle)**, and example iEEG contacts in a human clinical patient **(bottom)**. **(b)** Average power spectral density estimate for spindle windows detected by the CNN model (red) and matched non-spindle windows (black), illustrating the nearly 10 dB increase within the 11–15 Hz spindle band in NHP ECoG recordings **(top)**, human EEG recordings **(middle)**, and human iEEG recordings **(bottom)**. Power at line noise frequency omitted for clarity. **(c)** The architecture of the CNN model developed for spindle detection. **(d)** Examples of detected spindles by the CNN model (red) in NHP ECoG recordings **(top)**, human EEG recordings **(middle)**, and human iEEG recordings **(bottom)**.

What is the spatial extent of spindle oscillations across cortex? To answer this question, we studied the distribution of simultaneously detected spindles across recording sites. We defined three classes of spindles based on co-occurrence across recording sites: local (1-2 sites), regional (3-10 sites), and multi-area (more than 10 sites). We noted our CNN approach detected many spindles with electrode sites distributed widely across the cortex (Figure 2a). By taking the unique cortical regions covered by the electrodes into account, we verified these were multi-area spindle events (Figure 2b). We then compared spindles detected by the CNN and AT approaches. To do this, we computed the ratio of spindles detected by the CNN and AT for all classes. This comparison revealed multi-area spindles were systematically detected approximately 1.5 (ECoG) to 10 (iEEG) times more often with the CNN than with the AT (Figure 2c and Supplementary Figure 3). Across all recordings, the increase in the multi-area spindles detected by the CNN was significantly greater than in the local spindles (p < 1 x 10^−3^, NHP ECoG recordings; p < 1 x 10^−6^, EEG recordings; p < 0.02, iEEG recordings, one-sided Wilcoxon signed-rank test; similar results for the local-regional comparison, p < 0.01, EEG recordings; p < 0.02, iEEG recordings, one-sided Wilcoxon signed-rank test, n.s. in NHP ECoG). Taken together, these results demonstrate that spindles appear much more widespread across cortex when detected using our approximately amplitude-invariant deep learning approach.

**Figure 2.**
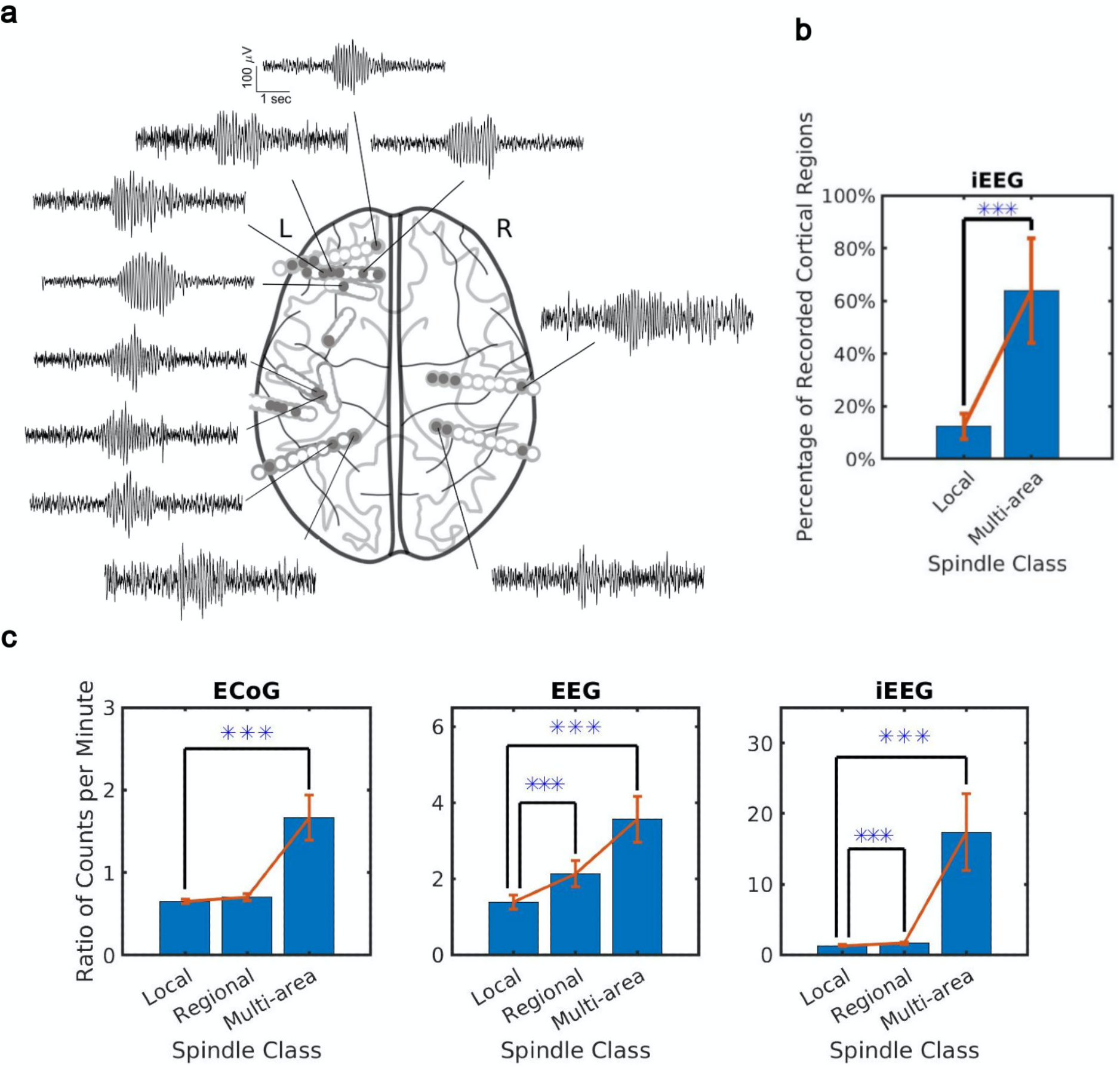
Distribution of the extent of spindles detected by CNN and AT approaches. **(a)** An example of a widespread, multi-area spindle with electrode sites distributed widely across the cortex. Filled gray circles indicate electrode contacts in gray matter. **(b)** Plotted is the percentage of unique recorded cortical regions with spindles detected by the CNN in the local versus multi-area case across all subjects in the iEEG recordings (average ± SEM). **(c)** Plotted are the ratios of spindles detected by the CNN and AT in NHP ECoG recordings **(left),** human EEG recordings **(middle)**, and iEEG recordings **(right)** in local (1-2 sites), regional (3-10 sites) and multi-area (more than 10 sites) spindle classes (average ± SEM in all cases). Across recordings, the increase in regional and multi-area spindles detected by the CNN is significantly larger than for the local spindles (except local vs. regional in the NHP ECoG).

The organization of spindles across the cortex is thus neither fully local nor fully global: the co-occurrence patterns of this sleep rhythm contain a mixture of local and widespread events. If this is the case, what is the impact of the distribution on sleep-dependent memory consolidation? To answer this question, we further studied the human EEG dataset, which had the unique feature of testing sleep after tasks with varying memory loads. Briefly, before nap EEG recordings, subjects completed a task in which five novel outdoor scenes (high visual working memory, H-WM) or two novel outdoor scenes (low visual working memory, L-WM) were required to be held in working memory for six seconds (Figure 3a). After the delay period, subjects were then presented with a subsequent visual scene and asked whether it belonged to the previously presented set. In each case (H-WM and L-WM), trials were balanced so that the same total number of visual scenes was presented before sleep. An increase in spindle density after memory tasks and its relationship with memory consolidation is well established (Clemens et al., 2005; Dang-Vu et al., 2008; Gais et al., 2002; Schabus et al., 2007, 2004); however, the effect of memory tasks on co-occurrence remains unknown. Considering the potential circuit mechanism for spindles to link activity in neuron groups distributed across multiple areas in cortex through long-range excitatory connections (Muller et al., 2016), we then hypothesized sleep following high-load visual memory tasks would exhibit more multi-area spindles and a larger spatial extent. To test this hypothesis, we first confirmed that amplitudes of detected spindles did not differ across L-WM and H-WM conditions (p > 0.77, Wilcoxon signed-rank test). We then computed the rate of multi-area spindles after L-WM and H-WM tasks. Both regional and multi-area spindles appeared more often after H-WM than L-WM (p < 0.032, regional spindles; p < 0.006, multi-area spindles; one-sided paired-sample t-test) as detected by the CNN model, consistent with our hypothesis (Figure 3b). Similarly, the largest increases following H-WM versus L-WM were observed in the subset of regional and multi-area spindles detected by the more-conservative SNR approach (Supplementary Figure 4a); however, no increase in multi-area spindles was observed with the AT algorithm (Supplementary Figure 4b).

**Figure 3.**
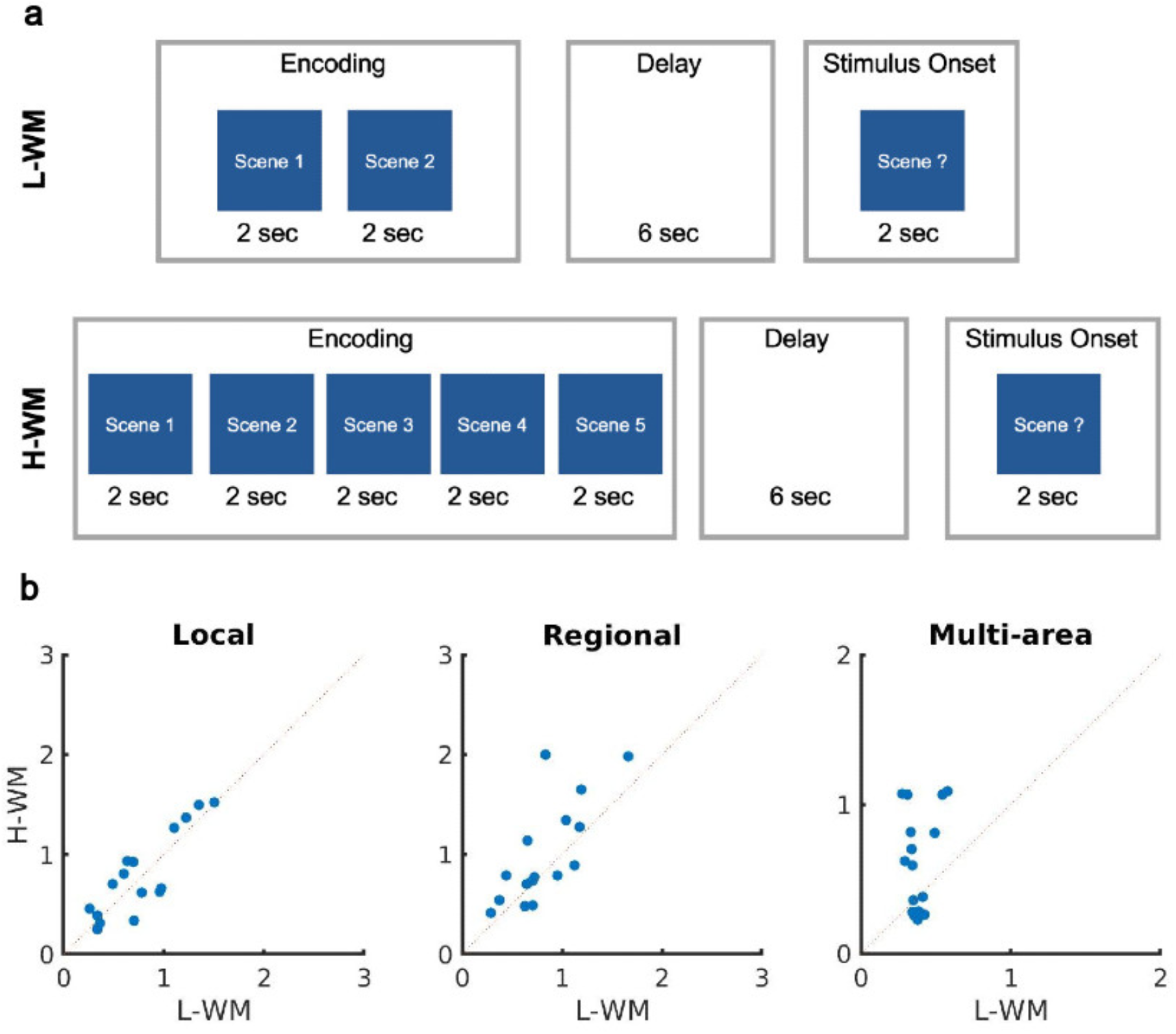
Impact of visual memory load on sleep spindle occurrence. **(a)** Schematic representation of low and high visual working memory tasks. **(b)** Average number of spindles detected per minute in high versus low visual working memory condition. Spindles are grouped into local **(left)**, regional **(middle)** and multi-area **(right)** classes detected by the CNN model. A significant increase in the number of spindles among subjects can be observed in multi-area and regional spindles as opposed to local spindles (p > 0.34, local spindles; p < 0.032, regional spindles; p < 0.006, multi-area spindles; one-sided Wilcoxon signed-rank test).

In this work, we have studied sleep spindles from human and NHP sleep recordings. To analyze these recordings, we adapted newly developed deep learning approaches for detecting rhythmic events in high-noise data (George and Huerta, 2018; Perol et al., 2018). We validated this approach through a series of control analyses and comparison to the subset of spindles detected by a similarly amplitude-invariant but more-conservative SNR algorithm. We found that large, multi-area spindles, where more than 10 electrode sites exhibit this rhythm simultaneously, are much more prevalent than previously estimated by amplitude-thresholding approaches (Andrillon et al., 2011; Nir et al., 2011; Piantoni et al., 2016; Sarasso et al., 2014), which tend to select only the highest-amplitude events.

While it has become increasingly clear that sleep spindles play an active and causal role in sleep-dependent memory consolidation (Aton et al., 2014; Clemens et al., 2005; Eschenko et al., 2006; Gais et al., 2002; Mednick et al., 2013; Rasch and Born, 2013), it remains unclear how these oscillations coordinate activity across areas to drive formation of strong neocortical assemblies distributed over long distances (Klinzing et al., 2019). Episodic memories often contain a detailed multisensory scene (Horner and Burgess, 2013; Tulving, 1983), and the activity patterns associated with these memories recruit distributed representations from association to primary sensory areas (Horner et al., 2015; Wheeler et al., 2000). In recent work, we found that sleep spindles can be organized into large-scale waves rotating across neocortex (Muller et al., 2016), and we hypothesized that these waves could provide a mechanism by which long-range excitatory connections between distant populations in cortex could be strengthened during memory consolidation in sleep. This potential mechanism for memory consolidation is interesting in light of recent research showing wide distribution of memory engrams (Kitamura et al., 2017; Roy et al., n.d.) and experience-dependent myelination formation in memory consolidation (Pan et al., 2020; Wang et al., 2020), both of which are consistent with the development of distributed assemblies across neocortex by these waves. In this work, by adapting deep learning algorithms for waveform detection in high-noise recordings (George and Huerta, 2018; Perol et al., 2018), we have found that large-scale, multi-area spindles occur much more often than previously reported. Following these observations, we then hypothesized that large, multi-area spindles exhibit an increase following high-load memory tasks. Consistent with this hypothesis, both the CNN and SNR methods detect an increase in multi-area spindles uniquely following high-load memory tasks. These results thus provide a specific neural mechanism by which memories can be stored in distributed neocortical networks during sleep.

## Methods

### Recordings

We study performance of the CNN model across three sleep datasets obtained from electrodes ranging from traditional scalp EEG to invasive intracranial depth electrodes. These datasets represent recordings from very different electrode types, which vary widely in resolution and signal-to-noise ratio. Training the CNN model in the same way over these very different recordings demonstrates the generality of the framework developed here; further, these results also represent a cross-species comparison of sleep-rhythm dynamics in NHP and human neocortex.

The first dataset contains electrocorticographic (ECoG) recording from most of the lateral cortex in two macaques during natural sleeping conditions (Yanagawa et al., 2013). Recordings were obtained from 128 electrodes in both monkeys and sampled at 1 kHz by a Cerebus data acquisition system (Blackrock Microsystems, Salt Lake City, UT, USA). The quality of sleep was studied by the degree of spatial synchronization in slow wave oscillations and significant increase in delta power was reported in sleep condition versus waking activity. This dataset was recorded and distributed by Laboratory for Adaptive Intelligence, BSI, RIKEN and was made freely available at http://neurotycho.org/sleep-task.

The second dataset contains high-density scalp electroencephalography (EEG) recording from 20 healthy participants (Mei et al., 2018). Each participant participated in two separate sessions and completed a high- and low-load visual working memory task. The recordings were obtained during naps following the working memory tasks from a 64-electrode EEG skull cap and sampled at 1 kHz. The recordings were reviewed and stage 2 sleep was manually annotated by an expert to verify quality of sleep recordings. Ultimately, sleep recordings that did not reach stage 2 sleep or were too noisy were excluded from the study. Under these criteria, four subjects were excluded (subject 12, 20, 26 and 27). In addition, the recordings were common average referenced (CAR) to remove large artifacts with potentially non-neural origin. These recordings were made freely available at the Open Science Framework through the link https://osf.io/chav7.

The last dataset contains intracranial electroencephalography (iEEG) recordings from 5 epileptic patients in the Epilepsy Monitoring Unit (EMU) at London Health Sciences Centre (LHSC). Patients were implanted using depth electrodes for the sole purpose of surgical evaluation. Informed consent was collected from the patients in accordance with local Research Ethics Board (REB) guidelines. Each patient was implanted with 9 to 14 iEEG electrodes located across the cortex with up to 10 contacts in gray or white matter (Supplementary Table 1). The iEEG signals were recorded continuously for a duration of 7 to 14 days for the purpose of seizure localization. We use clinically annotated sleep onsets and study half an hour recording starting from the beginning of the sleep/nap cycles in electrode contacts located within gray matter.

### Signal-to-noise ratio (SNR) measure for sleep spindle detection

To specify a subset of spindles required to train our convolutional neural network (CNN) model, we implemented a modified version of signal-to-noise ratio (SNR) algorithm (Muller et al., 2016). This algorithm, which is inspired by the adaptive, constant-false-alarm-rate (CFAR) technique in radar, was used to detect narrow-band rhythmic activities. We measure the ratio of power within the frequency band of interest (here, 9-18 Hz) to power in the rest of the spectrum (1-100 Hz bandpass, with band-stop at 9-18 Hz) at each electrode. The SNR measure is computed over a sliding window of time (500 ms) and produces an estimate of how power in the frequency band of interest compares to total power in the recording, taking into account the noise on individual electrodes. We then used the SNR algorithm to produce high-quality training samples for the CNN model. To do this, we reduced the probability of false positives by setting the threshold to the 99^th^ percentile of the SNR distribution, thus detecting only the activity patterns that have the highest unique power concentration in the spindle frequency range. We additionally required the SNR algorithm to only include activities with a duration between 0.5 to 3 seconds, consistent with the duration of sleep spindles. The detected windows are then used for training the CNN model.

To additionally verify performance of the SNR algorithm, we implement this approach over one second recordings of a 90 by 90 array of LFPs generated by a spiking network model of cortical activity in the awake state, which does not contain the thalamic reticular loops and thalamocortical projections needed to generate sleep spindles. SNR values calculated from these data were uniformly below 0 dB, confirming the robustness of our approach in uniquely detecting spindle-frequency activity through a known ground truth dataset.

### Convolutional neural network (CNN) for sleep spindle detection

We developed a convolutional neural network (CNN) to detect spindles activities during sleep. The model is motivated by the successful implementation of deep learning for detecting earthquakes and gravitational waves in high-noise settings (George and Huerta, 2018; Perol et al., 2018). If trained properly, it has the ability to detect hidden spindles that are being unnoticed and provides a great opportunity to study the spatial and temporal analysis of spindle activities across the cortex. We first tested CNN models with different architectures and selected one of the best architectures across the sleep recording datasets (Figure 1c). Our CNN model is a one dimensional model with 5 convolutional layers (with 32, 64, 128, 192 and 256 filters) and 4 fully connected layers (with sizes 128, 64, 32 and 2). Each convolutional layer is followed by a maxpool and rectified linear unit layers and the output of the fifth convolutional layer is gradually flattened into 2D vectors using the fully connected layers followed by rectified linear unit layers. Our classifier has an additional softmax layer at the end which returns the probability of spindle in addition to the predicted label. The CNN model takes a window of sleep recording (500 ms which is bandpass filtered at 1-100 Hz after removal of line noise and harmonics) as an input and predicts its label (spindle or non-spindle). The hyperparameters of the CNN model are optimized to minimize the difference in the predicted and actual labels determined by the SNR algorithm.

### Power-spectral density estimate

To verify performance of the CNN, SNR and AT approaches, we compared power spectral density (PSD) estimates of spindle and non-spindle activities (Welch’s method; Figure 1b and Supplementary Figure 5). In both cases, we first remove line noise artifacts. We then compute PSD over windows of 0.5 s with no overlap and average spectra over detected events. Matched non-spindle PSDs were estimated over a large number of randomly selected non-spindle windows. The increase in the power during the natural frequency range of sleep (~9-18 Hz) in spindle vs non-spindles activities demonstrates the ability of both the CNN model and SNR algorithm to correctly identify spindle activities.

### Time-shifted averaging control

As an additional control analysis, we computed average signals over detected spindles, with activity shifted to align the largest oscillation peak in the detected time window. To compute this average, we first needed to correct for the time offset between different spindles. To do this, we shifted detected spindles to the largest positive value within the detected window, corresponding to the positive potential of an individual spindle oscillation cycle, and then took the average over all time-shifted windows. The average of time-shifted signals is computed over spindle windows detected by the CNN approach, as well as matched randomly selected non-spindle windows. Importantly, while the time-shifted average clearly exhibits 11-15 Hz oscillatory structure when computed over spindle events detected by the CNN, this need not be the case, as demonstrated by application of the same approach to matched non-spindle events (Supplementary Figure 1).

We also systematically studied the sensitivity of the CNN model as a function of the SNR threshold used for building the training set. To do this, we computed the time-shifted average over spindle events detected by the CNN model at different levels of the SNR threshold (Supplementary Figure 6). Clear, well-formed 11-15 Hz oscillatory structure is observed in the time-shifted averages above 0 dB threshold, verifying the quality of detected spindles by the corresponding CNN models. However, the 11-15 Hz oscillatory structure starts to disappear below 0 dB because an SNR threshold below 0 dB introduces errors into the training sets by mislabeling noise signals as spindles. On the other hand, similar oscillatory shapes of time-shifted average above 0 dB confirms the ability of the CNN model to perform robustly while trained over different sets of clearly-formed spindles.

### Comparison with amplitude-threshold approach

The amplitude threshold (AT) approach has been used extensively in the literature to automatically detect spindles during sleep(Gais et al., 2002; Nir et al., 2011). In this approach, a spindle is detected when the amplitude of the bandpass signal stays above a threshold for a limited period of time (e.g. at least 500 ms; cf. Supplementary Figure 5 in Nir et al., 2011). To implement this approach, we first bandpass filter the signal at the frequency of 11-15 Hz and then compute the root mean square (RMS) of its amplitude over a sliding window of 0.5 seconds. A spindle is detected whenever the RMS amplitude stays above the predetermined threshold for 0.5 to 3 s. To determine the most appropriate threshold for comparison to the CNN and SNR approaches, we first computed the distribution of electrode-level RMS amplitude that results in approximately 2 spindles per minute and then set the overall threshold to its average across all electrodes. The quality and extent of detected spindles by the AT approach was then compared with the CNN and SNR (Figure 2c, Supplementary Figure 2, 3, 4, 5 and 7). The CNN model has a relatively amplitude-invariant nature in comparison with the amplitude-thresholding (AT) approach, which is highly sensitive to a predefined cutoff amplitude threshold. The AT approaches may only select spindles with the largest-amplitude events, or could miss ones that temporarily dip below the threshold while our approach has the ability to find well-formed spindles that are both large- and small-amplitude (Supplementary Figure 2).

### Electrode Localization

For the purpose of electrode localization in the iEEG recordings, we developed an image processing pipeline which involves electrode contact localization, brain tissue segmentation and atlas fitting. Semi-automatic contact localization was performed in 3D Slicer using the SEEG Assistant module (Narizzano et al., 2017). The entry and target points of each electrode were manually defined on the post-operative CT image. The entry/target labels were provided to the SEEGA algorithm, which automatically segmented the electrode contacts. To obtain brain location information for each contact, brain tissue segmentation and atlas fitting was carried out. To enable the use of anatomical priors during tissue segmentation, the pre-operative T1w MRI was non-linearly registered to the MNI152 2009c Nonlinear Symmetric template (https://www.bic.mni.mcgill.ca/ServicesAtlases/ICBM152NLin2009) using NiftyReg (Modat et al., 2010). An anatomical mask was generated by applying the inverse transform to the T1w image using the antsApplyTransforms algorithm from Advanced Normalization Tools 2.2.0 (ANTS; http://stnava.github.io/ANTs). Segmentation of gray matter, white matter, and cerebrospinal fluid was performed using the Atropos algorithm from ANTS(Avants et al., 2011b), which implements k-means classification (k=3). The resulting posteriors were merged into a 4D volume using the fslmerge algorithm from FMRIB Software Library v6.0 (FSL; https://fsl.fmrib.ox.ac.uk/fsl/fslwiki). The CerebrA atlas (Manera et al., 2020) was used to obtain anatomical labels for each electrode contact. Normalization to template space (MNI152NLin2009cAsym) was performed using the non-linear SyN (Avants et al., 2011a) symmetric diffeomorphic image registration algorithm from ANTS, using both the brain masks of the pre-operative T1w and template space. Using the inverse of the non-linear transform, the CerebraA atlas labels were warped to the pre-operative T1w MRI space. The atlas labels were then dilated using the fslmaths algorithm from FSL. The final T1w brain tissue/atlas segmentation was mapped to the contacts to provide location information for each contact (tissue probability and brain anatomical region). This custom processing pipeline has been made available on GitHub (https://github.com/akhanf/clinical-atlasreg).

## Code availability

Our custom MATLAB (MathWorks) implementations of all computational analyses, along with the analysis scripts used for this study will be made available as an open-access release on GitHub (http://mullerlab.github.io).

## Supplementary Figures

**Supplementary Figure 1.**
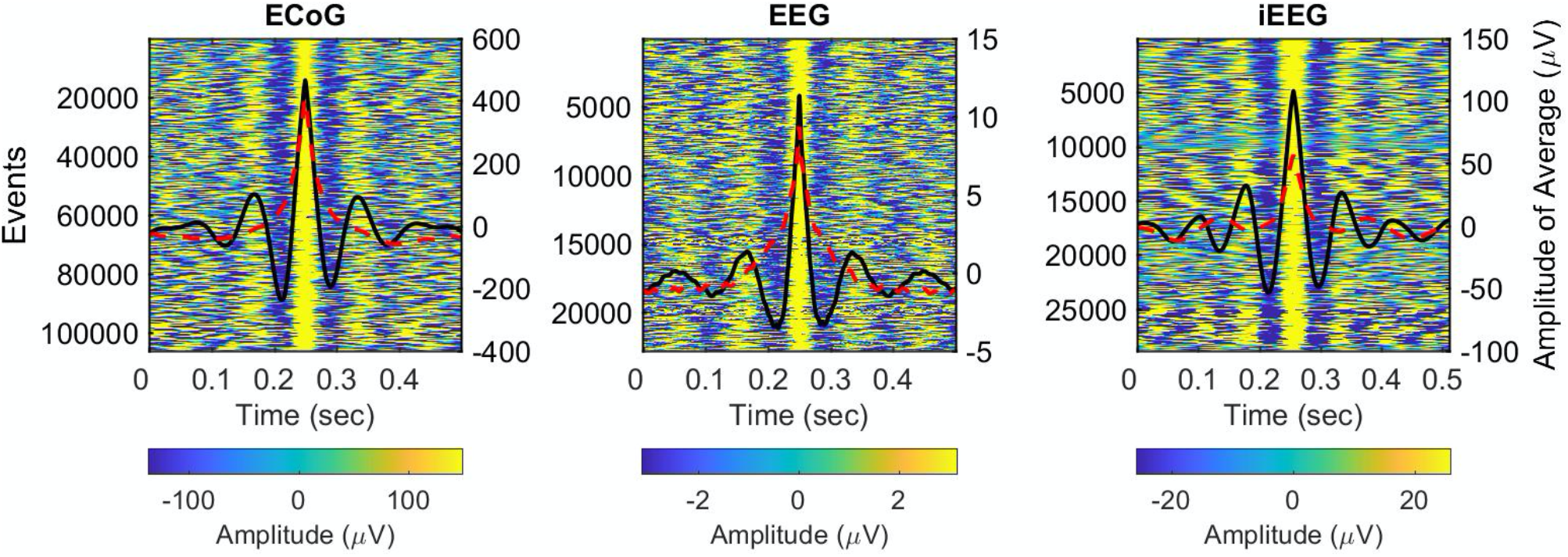
Average time-shifted spindles detected by the CNN model. Examples of average signals computed over spindle windows detected by the CNN models (solid black lines) vs a subset of randomly matched non-spindle windows (dashed red lines) in NHP ECoG recordings **(left)**, human EEG recordings **(middle)**, and iEEG recordings **(right)**. The average over detected spindle windows exhibits clear 11-15 Hz oscillatory structure, while no oscillatory structure is present in the average over matched non-spindle windows. The heatmaps in the background are the individual time-shifted spindle events, which demonstrate the presence of this structure in individual instances.

**Supplementary Figure 2.**
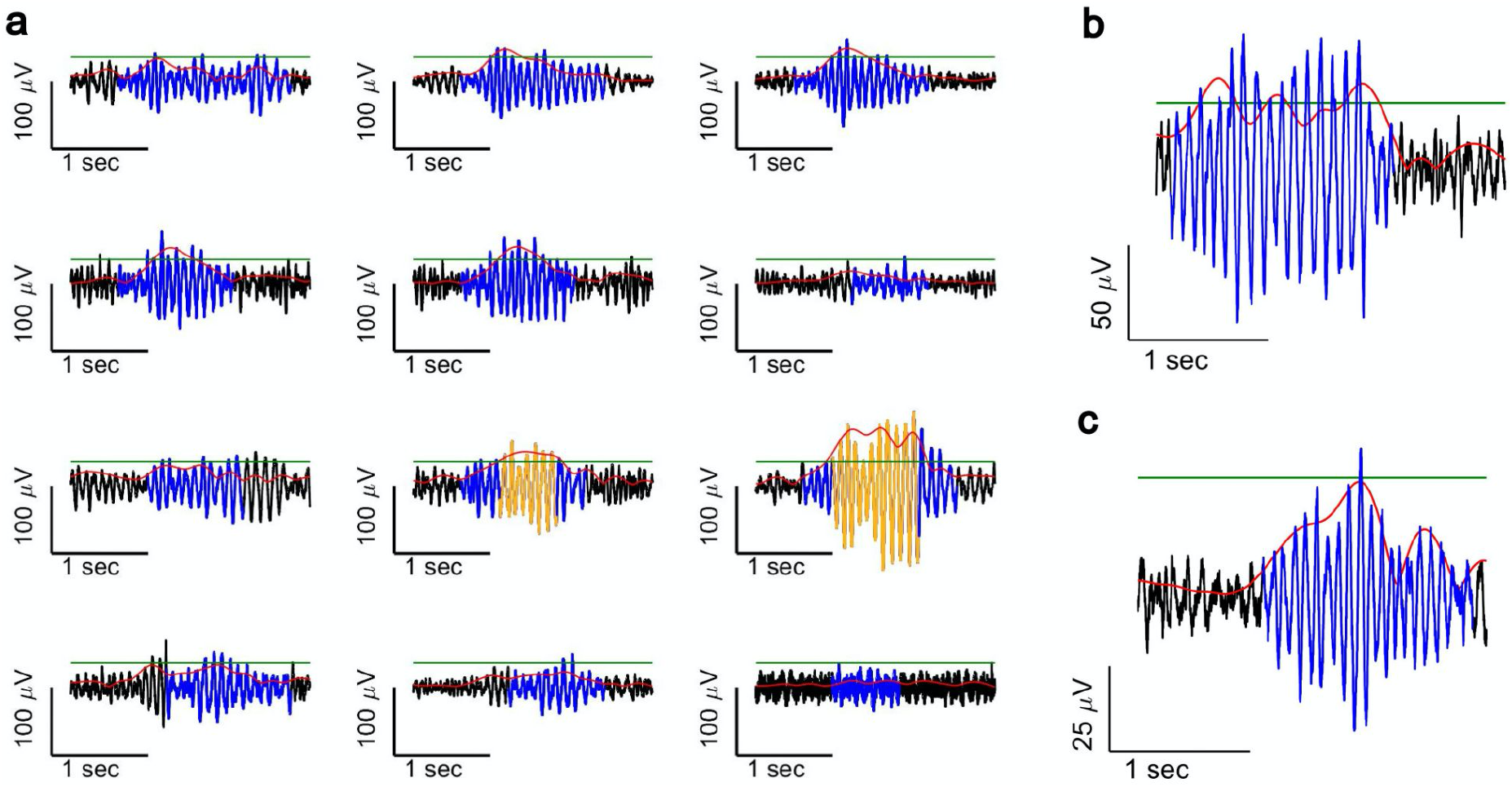
Performance of CNN model versus AT Algorithm. **(a)** An example of co-occurring spindles detected by the CNN model in 11 sites. The red line is the root mean square amplitude of the signal, the green line is the amplitude threshold, and the blue line represents windows of time in which spindles were detected by the CNN model. The AT algorithm, which only detects spindle activities with amplitude above a predefined amplitude threshold for at least a duration of 0.5 sec (Nir et al., 2011) (yellow line), was not successful in detecting the majority of spindles in this example. Specifically, the AT algorithm fails to detect well-shaped spindles (detected by the CNN model) whose amplitudes temporarily drop below threshold **(b)** or have low amplitude **(c).**

**Supplementary Figure 3.**
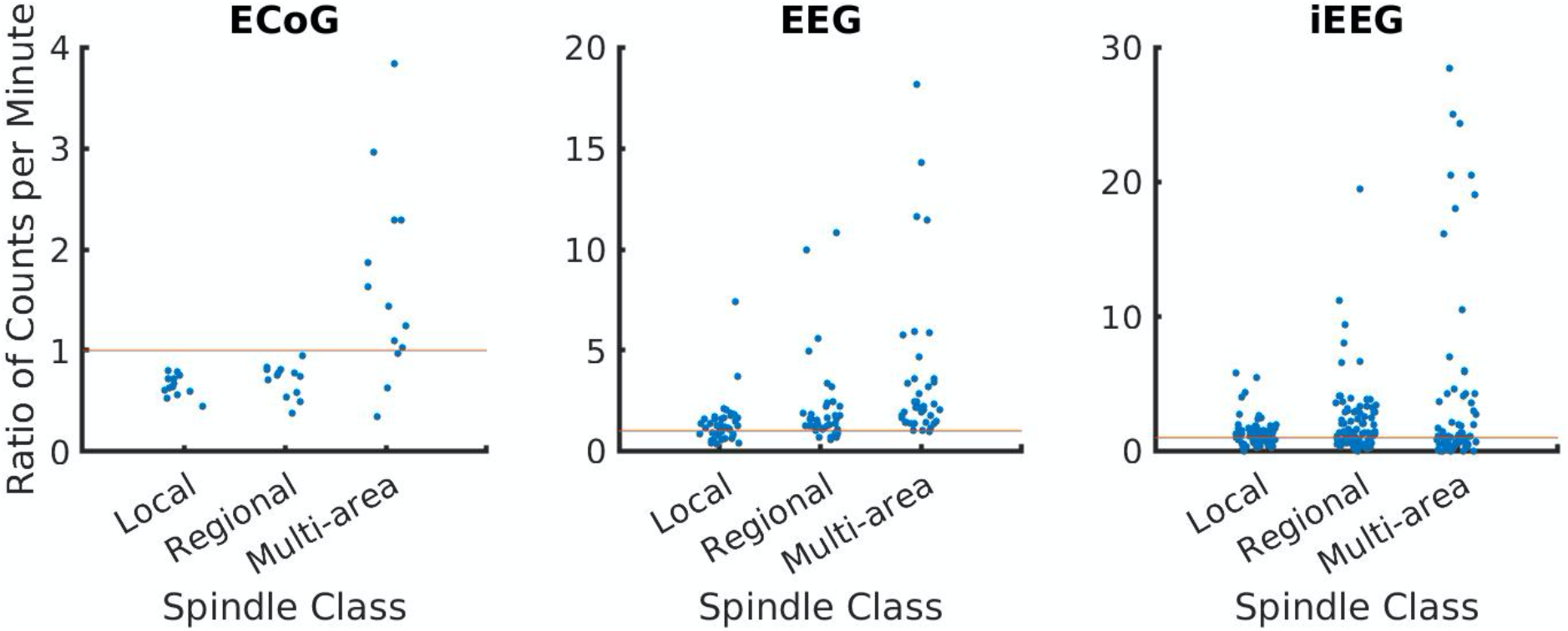
Distribution of the extent of spindles detected by CNN and AT approaches. Plotted is the scatter diagram of the ratio of average number of detected spindles by the CNN and AT in NHP ECoG recordings **(left),** human EEG recordings **(middle)**, and iEEG recordings **(right)** grouped into local (1-2 sites), regional (3-10 sites), and multi-area (more than 10 sites) spindle classes. Each blue dot represents the ratio of the number of spindles per minute in one sleep recording.

**Supplementary Figure 4.**
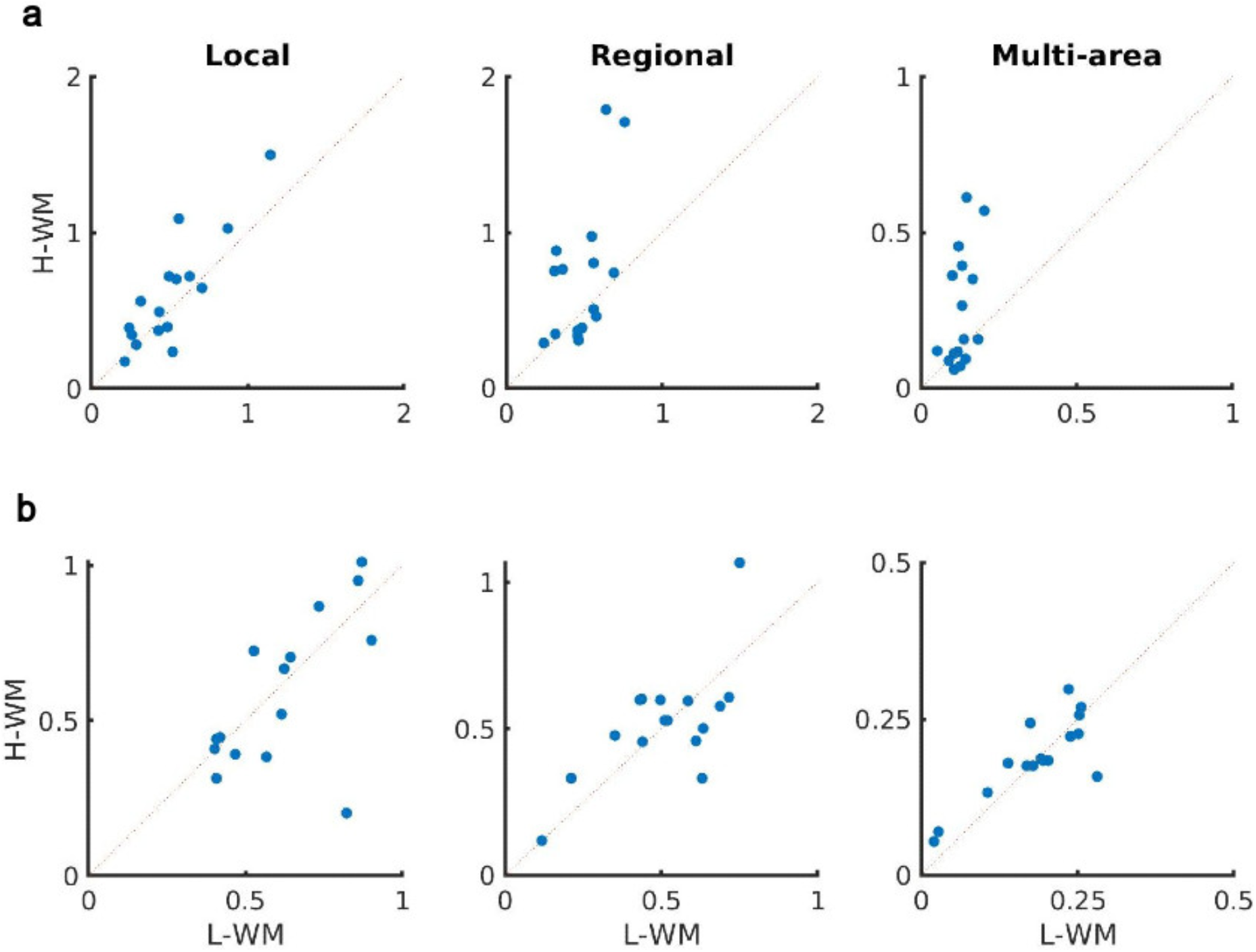
Low and high visual memory task and its impact on sleep spindle occurrence - SNR and AT. **(a)** Average number of spindles detected per minute in high vs low visual working memory condition by the SNR. The SNR algorithm detected a significant increase in the number of spindles across subjects (p < 0.04, local spindles; p < 0.02, regional spindles; p < 0.007, multi-area spindles; one-sided paired-sample t-test), with the largest increases for regional and multi-area spindles. **(b)** Average number of spindles detected per minute in high vs low visual working memory condition by the AT. In contrast to the CNN, AT was not able to detect any significant increase in the number of distributed spindles among subjects across all spindles classes (p > 0.56, local spindles; p > 0.34, regional spindles; p > 0.30, multi-area spindles; one-sided paired-sample t-test).

**Supplementary Figure 5.**
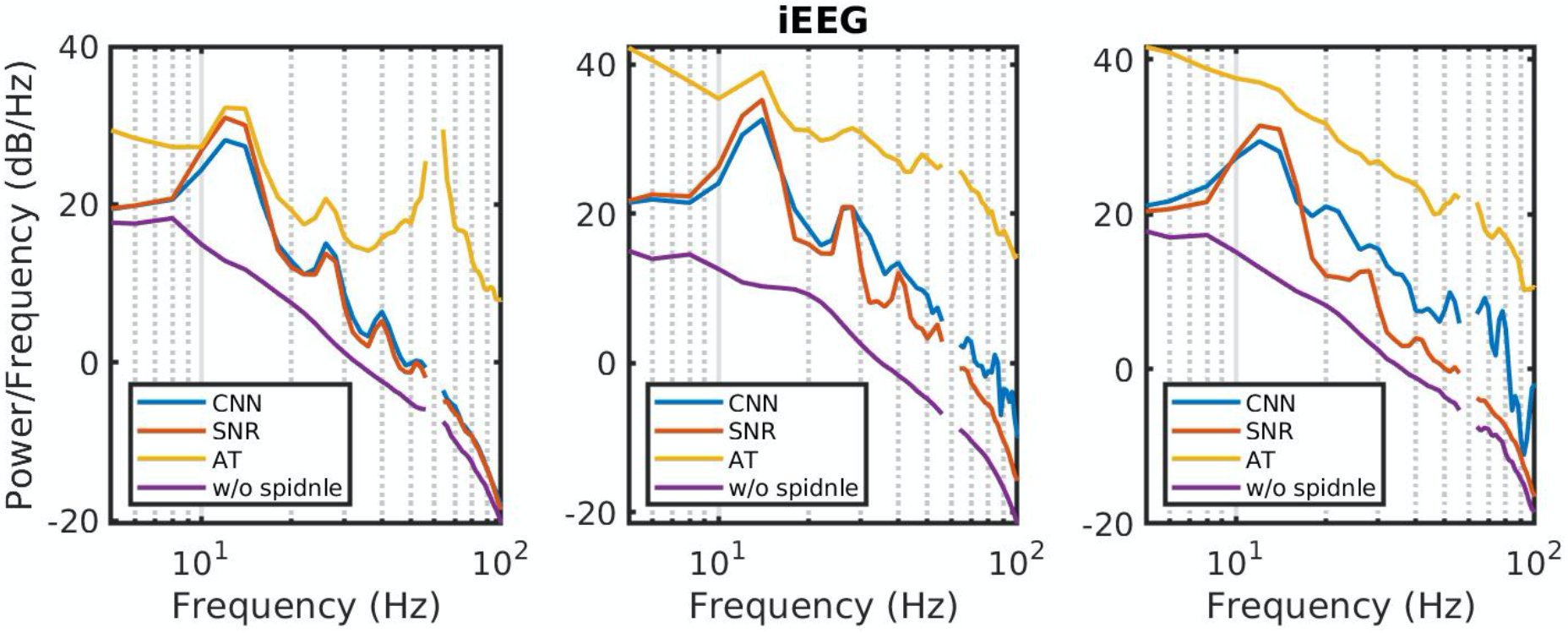
Power spectral density (PSD) comparison. Examples of average power spectral density estimate over spindle windows detected by the CNN model (blue), the SNR approach (red), the AT algorithm (yellow) and matched non-spindle windows (purple) in iEEG recordings. The CNN and SNR PSDs exhibit a nearly 10 dB increase within the 11–15 Hz spindle band compared to matched non-spindle windows, while the AT PSD exhibits higher power outside of the frequency of interest in addition to spindle band. Power at line noise frequency omitted for clarity.

**Supplementary Figure 6.**
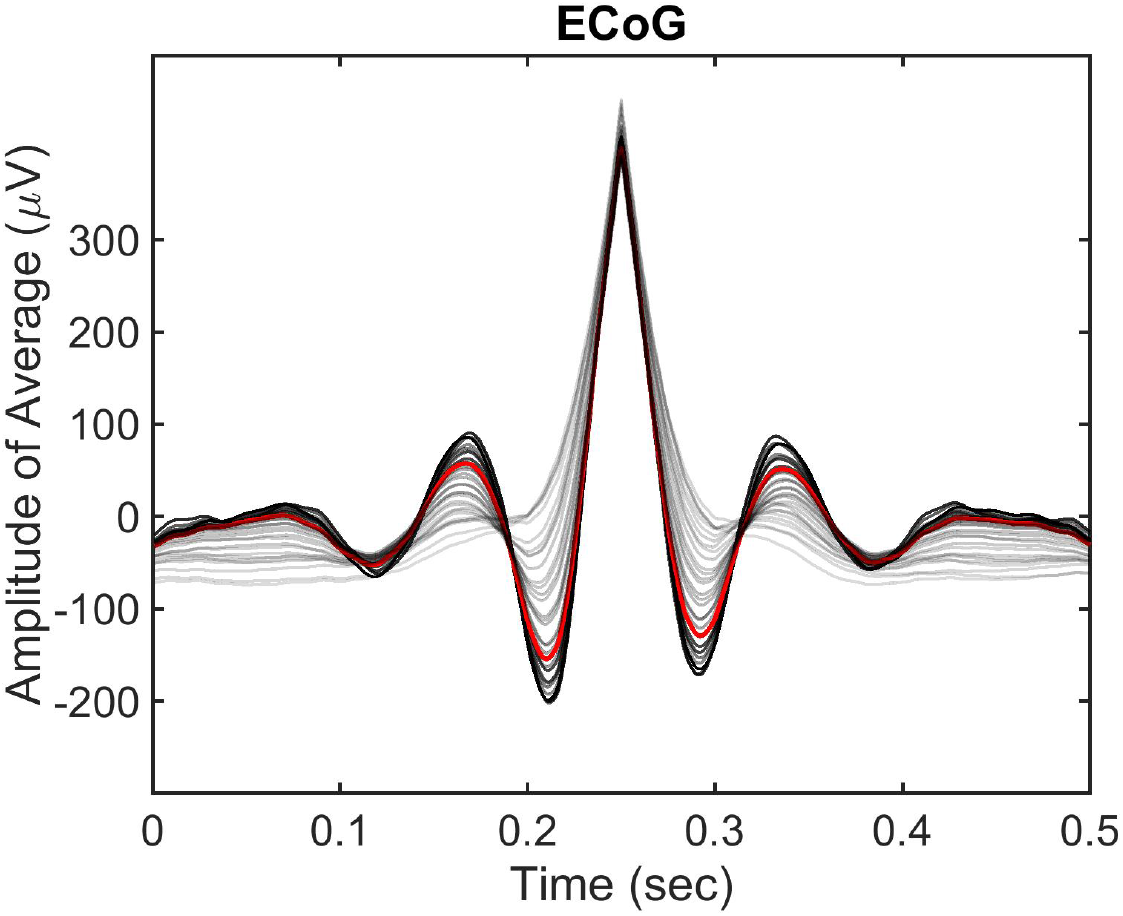
Impact of SNR threshold on the CNN model. Plotted is the change in the average time-shifted spindle detected by CNN models trained over a wide range of SNR thresholds (5 to −10 dB, dark to light gray) in the NHP ECoG recordings. Clearly detected spindle activity decreases with the SNR threshold, demonstrating that the CNN result breaks down when the model is trained on lower quality examples. The red line represents the average computed at 0 dB threshold (which represents parity between power in the spindle passband and the rest of the signal spectrum), below which the average detected spindle activities starts to drift away from the expected 11-15 Hz oscillatory structure.

**Supplementary Figure 7.**
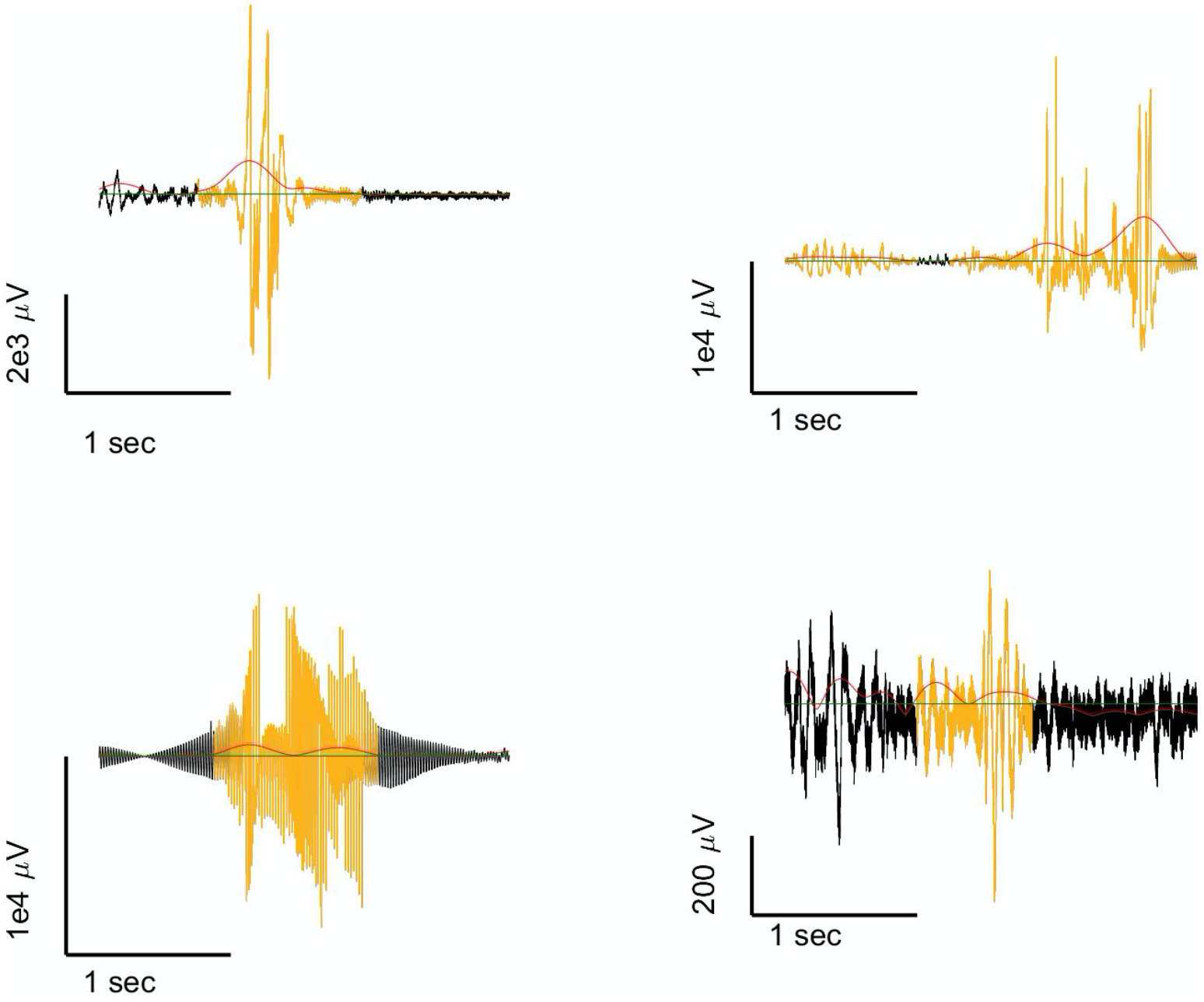
Non-spindle activities detected by the AT algorithm. Sleep recordings are subjected to artifacts, such as line noise, electrical noise, and movement artifacts that introduce signal distortion. These artifacts can result in false spindle activity detection in the AT approach. For example, the sharp artifact when filtered at 11–15 Hz in the AT approach appears as an oscillation which does not exist in the original recording **(top row),** or the AT approach might detect non-spindle activities resulting from broken channels **(bottom left)** or not clearly formed spindles **(bottom right)**.

---

**Supplementary Table 1:** Distribution of gray matter contacts in the cortical regions of all subjects in the iEEG recordings.

| Brain Region | Subject A | Subject B | Subject C | Subject D | Subject E |
| --- | --- | --- | --- | --- | --- |
| Amygdala | 0 | 5 | 4 | 4 | 0 |
| Caudal Middle Frontal | 0 | 0 | 0 | 0 | 3 |
| Entorhinal | 0 | 1 | 0 | 0 | 0 |
| Fusiform | 0 | 1 | 0 | 3 | 0 |
| Hippocampus | 8 | 12 | 8 | 6 | 0 |
| Inferior Lateral Ventricle | 0 | 0 | 2 | 0 | 0 |
| Insula | 1 | 27 | 17 | 1 | 9 |
| Lateral Orbitofrontal | 0 | 3 | 2 | 0 | 6 |
| Lateral Ventricle | 1 | 1 | 0 | 0 | 0 |
| Medial Orbitofrontal | 1 | 1 | 1 | 0 | 2 |
| Middle Temporal | 3 | 4 | 8 | 9 | 0 |
| Pars Opercularis | 0 | 0 | 1 | 0 | 0 |
| Pars Orbitalis | 0 | 1 | 0 | 0 | 0 |
| Pars Triangularis | 1 | 1 | 2 | 0 | 4 |
| Rostral Anterior Cingulate | 1 | 2 | 1 | 0 | 4 |
| Rostral Middle Frontal | 7 | 2 | 6 | 0 | 9 |
| Superior Frontal | 1 | 0 | 4 | 0 | 17 |
| Superior Temporal | 4 | 0 | 0 | 2 | 0 |
| Transverse Temporal | 0 | 0 | 0 | 1 | 0 |

